# Petascale Homology Search for Structure Prediction

**DOI:** 10.1101/2023.07.10.548308

**Authors:** Sewon Lee, Gyuri Kim, Eli Levy Karin, Milot Mirdita, Sukhwan Park, Rayan Chikhi, Artem Babaian, Andriy Kryshtafovych, Martin Steinegger

## Abstract

The recent CASP15 competition highlighted the critical role of multiple sequence alignments (MSAs) in protein structure prediction, as demonstrated by the success of the top AlphaFold2-based prediction methods. To push the boundaries of MSA utilization, we conducted a petabase-scale search of the Sequence Read Archive (SRA), resulting in gigabytes of aligned homologs for CASP15 targets. These were merged with default MSAs produced by ColabFold-search and provided to ColabFold-predict. By using SRA data, we achieved highly accurate predictions (GDT_TS > 70) for 66% of the non-easy targets, whereas using ColabFold-search default MSAs scored highly in only 52%. Next, we tested the effect of deep homology search and ColabFold’s advanced features, such as more recycles, on prediction accuracy. While SRA homologs were most significant for improving ColabFold’s CASP15 ranking from 11th to 3rd place, other strategies contributed too. We analyze these in the context of existing strategies to improve prediction.

## Introduction

Determining the 3D structure of proteins is of great importance to many research fields, encompassing cancer drug discovery (Ren et al. 2023; Borkakoti and Thornton 2023), pesticide development, and crop improvement (Koesoema 2022). It also plays a crucial role in the design of sensors and enzymes (Pereira et al. 2021), as well as numerous other applications, as reviewed by Pearce and Zhang (2021).

Traditionally, protein structures have been solved using laborious techniques, such as X-ray crystallography, resulting in just under 200,000 structures in over 50 years of communal effort (Berman et al. 2000; Subramaniam and Kleywegt 2022). Resolved structures are routinely deposited in the Protein Data Bank (wwPDB consortium 2019). The demanding experimental process has motivated the development of computational tools as a less burdensome alternative for structure prediction. Since 1994, the Critical Assessment of protein Structure Prediction (CASP) has aimed to identify state-of-the-art computational methods by competition (Moult et al. 1995). The organizers of CASP provide the participants with protein sequences, whose structures were experimentally solved but not yet deposited in the PDB. The solved structures are unknown to the organizers, assessors as well as to the participants. Also, the group identities are kept anonymous from the assessors, therefore the competition is considered double-blinded, ensuring fairness.

Originally, computational prediction methods could be divided into two main groups: template-based modeling (TBM) and free modeling (FM). However, in the past decade, the lines between the groups have been blurred (Bertoline et al. 2023). TBM is a broad category in which known structures are used as templates to predict the structure of query proteins, based on the sequence similarity between them. In its simplest form, a similarity between a single query and a match from the PDB serves as the base for projecting the match’s structure onto the query. The inaugural software MODELLER (Sali and Blundell 1993) and other tools that followed have made use of this principle -see Pearce and Zhang (2021) for a review.

Given the limited size of the PDB and its bias towards model organisms (Orlando et al. 2016), detecting remote sequence homology is crucial. To that end, increasingly sensitive search methods have been developed. The first step forward was taken by algorithms like BLAST (Altschul et al. 1990), which directly compare the query sequence to the reference database. PSI-BLAST (Altschul et al. 1997) improved upon this by computing a multiple sequence alignment (MSA) of the query and its best BLAST hits and calculating a position-specific scoring matrix from that. This generalization of the query is used for a sensitive search of the reference. This approach was further refined by using probabilistic hidden Markov models (HMMs) (Krogh et al. 1994) in tools like HMMer (Eddy 2011). Another significant advancement came with HHsearch (Söding 2005), which expressed both query and reference as HMMs, markedly improving search sensitivity. This underpinned the success of HHpred (Hildebrand et al. 2009) in the CASP9 challenge (Moult et al. 2011). A further development, HHblits (Remmert et al. 2011; Steinegger et al. 2019a) accelerated the HMM-HMM comparison allowing to query databases with millions of HMMs like the Uniclust30 (Mirdita et al. 2017), a clustered version of the Uniprot (UniProt Consortium 2023) to generate diverse query MSAs.

Due to its unprecedented sensitivity, HHpred has transformed CASP in two ways. First, many methods competing in CASP have incorporated HHblits/HHsearch or other tools to identify distant structural homologs (Bertoline et al. 2023). Second, CASP has started using it for classifying target domains in subsequent competitions (Kinch et al. 2011). Specifically, domains in targets for which HHpred could identify a homolog in the PDB, are considered by CASP as “TBM target domains”, while the others -as “FM target domains”.

Recent advances in deep learning have been harnessed by various methods for protein structure prediction (Torrisi et al. 2020). Undoubtedly, the most revolutionary of these is AlphaFold2 (Jumper et al. 2021), which won the CASP14 challenge by a significant margin (Kryshtafovych et al. 2021), reaching experimental accuracy for over two-thirds of the targets. Despite its success, AlphaFold2’s prediction accuracy is not without its limitations. Most notably, it relies on its input MSA (Mirdita et al. 2022) diversity, experiencing a significant drop in prediction accuracy when the median number of diverse sequences in the MSA is 30 or less (Jumper et al. 2021). This finding is in agreement with previous studies on the importance of distant homologs to structure prediction (Kuhlman and Bradley 2019; Ashkenazy et al. 2009). However, the ability to construct a deep MSA depends not only on the sensitivity of search algorithms, such as HHblits, but also on the potential pool of sequences, i.e., the reference database.

Metagenomics allows for sequencing uncultivable organisms directly from the environment, significantly expanding the repertoire of protein sequences deposited in scientific databases. In recent years, metagenomic sequences have shown great potential in increasing the fraction of proteins, whose structure can be modeled accurately (Söding 2017; Ovchinnikov et al. 2017; Yang et al. 2021; Wang et al. 2019). Of note, the largest metagenomic database used in these studies is the IMG/M, which contains 27 tera base pairs (Chen et al. 2023b).

It is therefore not surprising that the top scoring servers in the most recent CASP15 challenge were based on AlphaFold2 and included metagenomic sequences in their constructed MSAs (Table 1). An example for such a server is ColabFold (Mirdita et al. 2022). ColabFold takes as input a query protein sequence(s), whose structure is to be predicted. Its first step, denoted here as CF-search, implements a procedure for collecting homologs of the query using MMseqs2 (Steinegger and Söding 2017). CF-search starts by querying the input against the UniRef30 database (Mirdita et al. 2017) and computing profiles from the hits. Next, CF-search queries these profiles against one of two metagenomic databases, which were constructed as part of the ColabFold release: BFD/MGnify and ColabFoldDB (the default reference database). As detailed in Table 2, BFD/MGnify contains 513 million non-redundant proteins from the union of the BFD (Jumper et al. 2021) and MGnify (Richardson et al. 2023) databases. ColabFoldDB expanded the BFD/MGnify with various environmental proteins, resulting in ∼740 million proteins. Following the search, an MSA is computed from the detected homologs and finally, in a step denoted here as CF-predict, the MSA is provided as input to the AlphaFold2 models.

**Table 1.** Use of homology algorithms and databases among leading CASP15 servers^a^.

| **Name** | **Rank<sup>b</sup>** | **Homology search alg.** | **Ref. sequence DBs<sup>c</sup>** |
| --- | --- | --- | --- |
| Yang-Server | 1 | HHblits, MMseqs2, manual? | MGnify, UniRef30 |
| UM-TBM | 2 | DeepMSA, LOMETS | BFD, IMG/M, Metaclust, MetaSource, Uniclust30, UniRef90, Tara? |
| Manifold-E | 3 | HHblits, hmmsearch, Jackhmmer | BFD, Uniprot, UniRef30, UniRef90 |
| DFolding | 4 | CRFalign, Jackhmmer, HHblits, HHpred, Kalign | BFD, Uniclust30, UniRef90, MGnify |
| MULTICOM | 5-7,9 | DeepMSA (modified), Foldseek, HHblits, Jackhmmer, MMseqs2 | BFD, ColabFoldDB, IMG/M, MGnify, Uniclust30, UniRef90, Uniprot |
| RaptorX | 8 | HHblits, Jackhmmer | BFD, SMAG+MetaEuk+TOPAZ+MGV+GPD+IMG/M (in-house HHblits DB), MGnify, Uniclust30, UniRef90 |
| MultiFOLD | 10 | MMseqs2, UniRef30 | ColabFoldDB |
| ColabFold | 11 | MMseqs2 | ColabFoldDB, UniRef30 |

<sup>a</sup> The information about the servers was mined from the CASP15 abstract book.
<sup>b</sup> The rank refers to the "server only" performance on protein targets, excluding the TBM-easy category.
<sup>c</sup> A detailed overview of the reference sequence DBs is provided in Table 2.

**Table 2.**
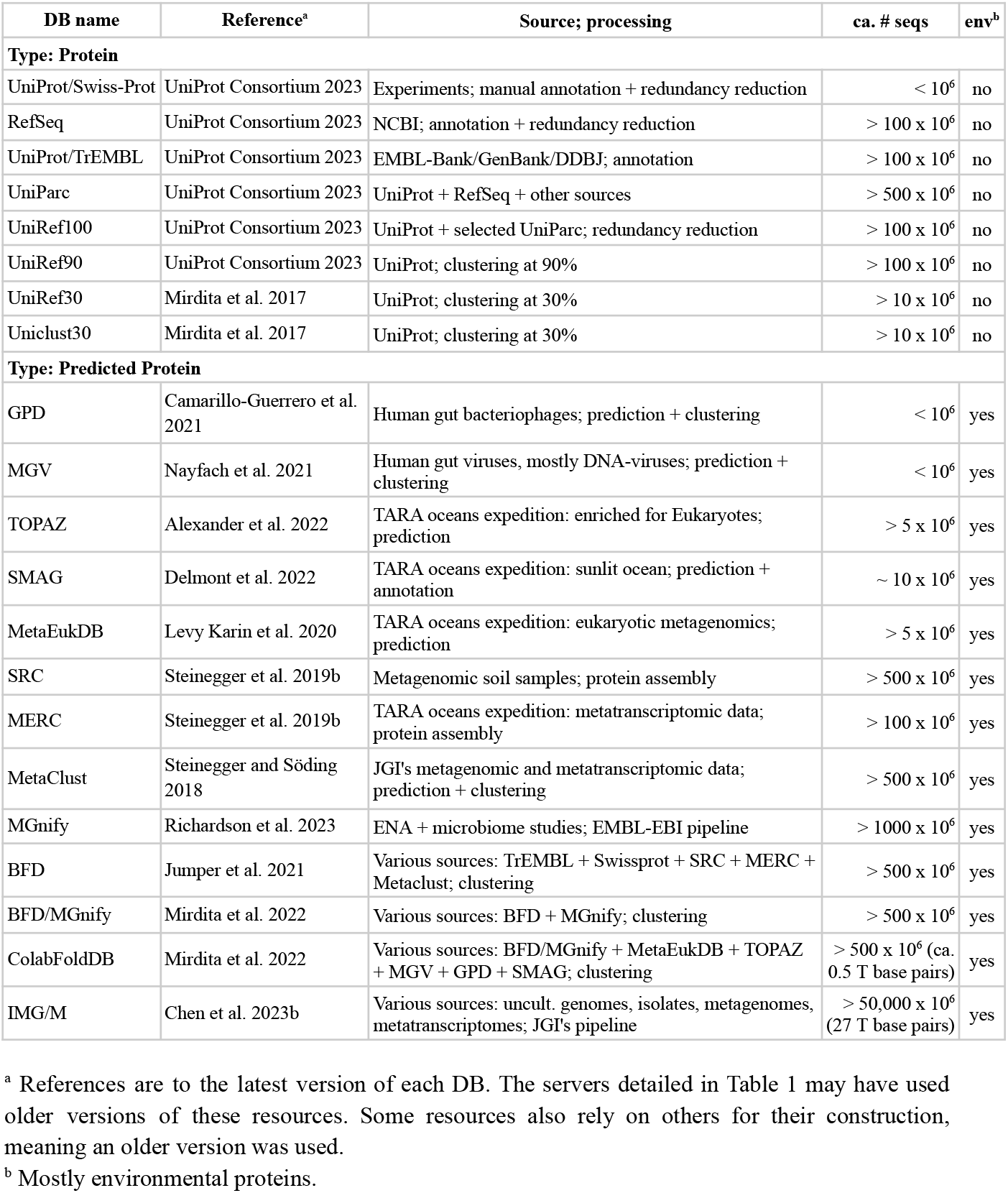
Size and composition of reference databases used by leading CASP15 servers.

In this study we examined different strategies to improve protein structure prediction along three axes. The first two focused on adding homologs to MSAs used for protein structure prediction and the third -on utilizing advanced features of CF-predict. The first and main axis is the breadth of the search, where we studied the impact of a much more systematic inclusion of metagenomic sequences on prediction accuracy. Over 37 peta base pairs are publicly available through the Sequence Read Archive (SRA), the world’s largest metagenomic database (Katz et al. 2022). Recently, Serratus, a tool for a high-throughput search of the SRA, was introduced (Edgar et al. 2022). Here, we constructed MSAs based on Serratus-mined homologs of the CASP15 targets and merged them with the default MSAs produced by CF-search. In the second axis of this study, we further enhanced the merged MSAs by searching for distant homologs of their sequences using HHblits against the BFD. The third axis concerns tuning the advanced parameters of ColabFold to control the use of templates and multimer models and the number of recycles. We then provided MSAs produced by each of these strategies as input to CF-predict and compared the resulting prediction accuracy to that measured with default CF-search MSAs as well as to the leading CASP15 servers. For fair comparison, we ensured all databases used in this study excluded any sequences deposited after the start of CASP15 competition (May 2022).

Our results show that adding SRA-mined homologs improves prediction accuracy for 61% of the examined targets. Tuning advanced features of CF-predict, especially adding more recycles, also contributed to better prediction. By combining the different strategies, ColabFold’s CASP15 ranking among servers on non-easy template targets increased from 11th to 3rd place, indicating the vast potential of large-scale sequence exploration for better structure prediction.

## Results and Discussion

### 1. Homolog search and MSA construction

The entry point to this study was a list of 126 targets provided by CASP15. We excluded from this list all targets, which were RNA, heteromers, canceled by CASP or indicated as auxiliary structure for ligand prediction, leaving 77 targets. Each of these targets had one or more domains, which are divided into categories by CASP as follows: FM, FM/TBM, TBM-hard and TBM-easy. We used CF-search to query the targets against ColabFoldDB (Fig. 1 ➀), resulting in 77 MSAs, denoted here as *cfdb* MSAs.

**Figure 1.**
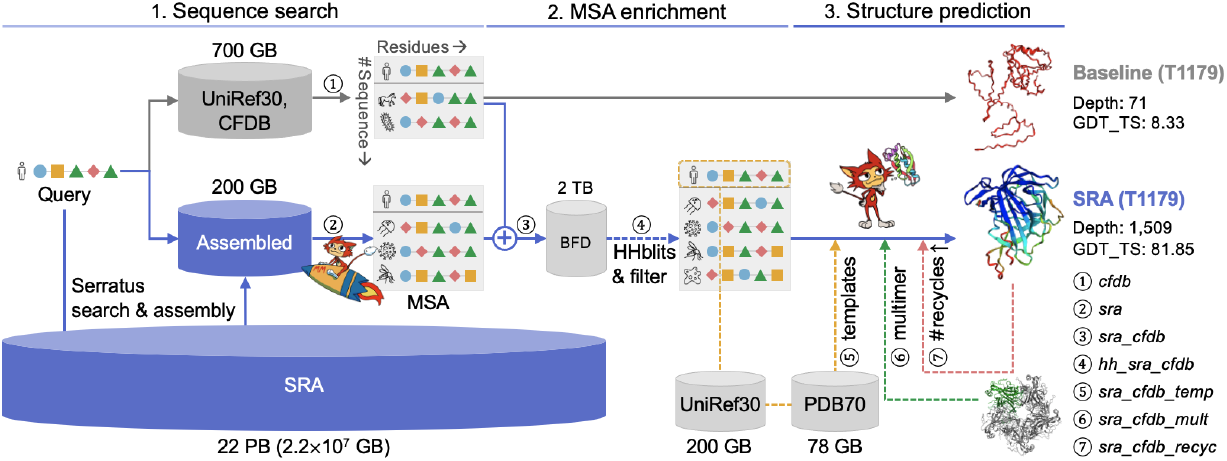
MSA enrichment using SRA and other strategies to improve protein structure prediction. Workflow of the different strategies examined in this study ➀∼➆. All strategies construct an MSA (but differ in the homology DBs they utilize) and provide it to CF-predict (but differ in the way they tune its parameters). The size of each homology DB is denoted close to it. The baseline MSA (*cfdb* MSA, ➀) is constructed by CF-search. The SRA-detected homologs are aligned to create ➁ using MMseqs2. The *sra_cfdb* MSA (➂) is constructed by combining ➀ and ➁. The *hh_sra_cfdb* MSA (➃) is constructed by querying ➂against UniRef30 and BFD using HHblits. Strategies ➄, ➅, ➆ refer to the following CF-predict options: use of templates, multimer (homo-oligomer) modeling, and 12 recycles (instead of the default 3). Before being provided to CF-predict, each MSA is filtered based on the sequence identity between its members and the query.

#### 1.1 Thousand times broader search

Our next goal was to expand the search beyond ColabFoldDB and explore the SRA. With its 37 peta base pairs of publicly available data, the SRA is orders of magnitude bigger than any previously-used metagenomic resource, including ColabFoldDB (Table 2). We queried the 77 CASP15 targets using Serratus against over half of the publicly available SRA, comprising 22 peta base pairs, organized in over five million SRA runs. Reads aligned to the CASP15 protein sequence queries, were examined in the context of their run and assembled using rnaviralSpades (Meleshko et al. 2021), mounting to over a hundred gigabytes of assembled data.

Each CASP15 target was then queried against a reference protein database created from its Serratus-produced assembled proteins using MMseqs2 (Steinegger and Söding 2017). The identified homologs were then aligned by MMseqs2 to create an MSA (Fig. 1 ➁), denoted as *sra* MSA. As a first indication of the tremendous capacity of the SRA, we found that more than half of the targets, which were composed solely of TBM-easy domains, could be matched with at least 1,000,000 homologs, some even exceeding 100,000,000 (Fig. 2A). Processing MSAs with millions of sequences poses a heavy computational burden. Thus, we opted to exclude from this study targets, which only contained TBM-easy domains, focusing on the remaining 46 targets, which had 62 non-TBM-easy domains: 39 FM, 8 FM/TBM, and 15 TBM-hard. On average, the number of homologs detected per domain doubled from 106,586 in the CFDB to 274,231 when including the SRA results (Fig. 2A).

**Figure 2.**
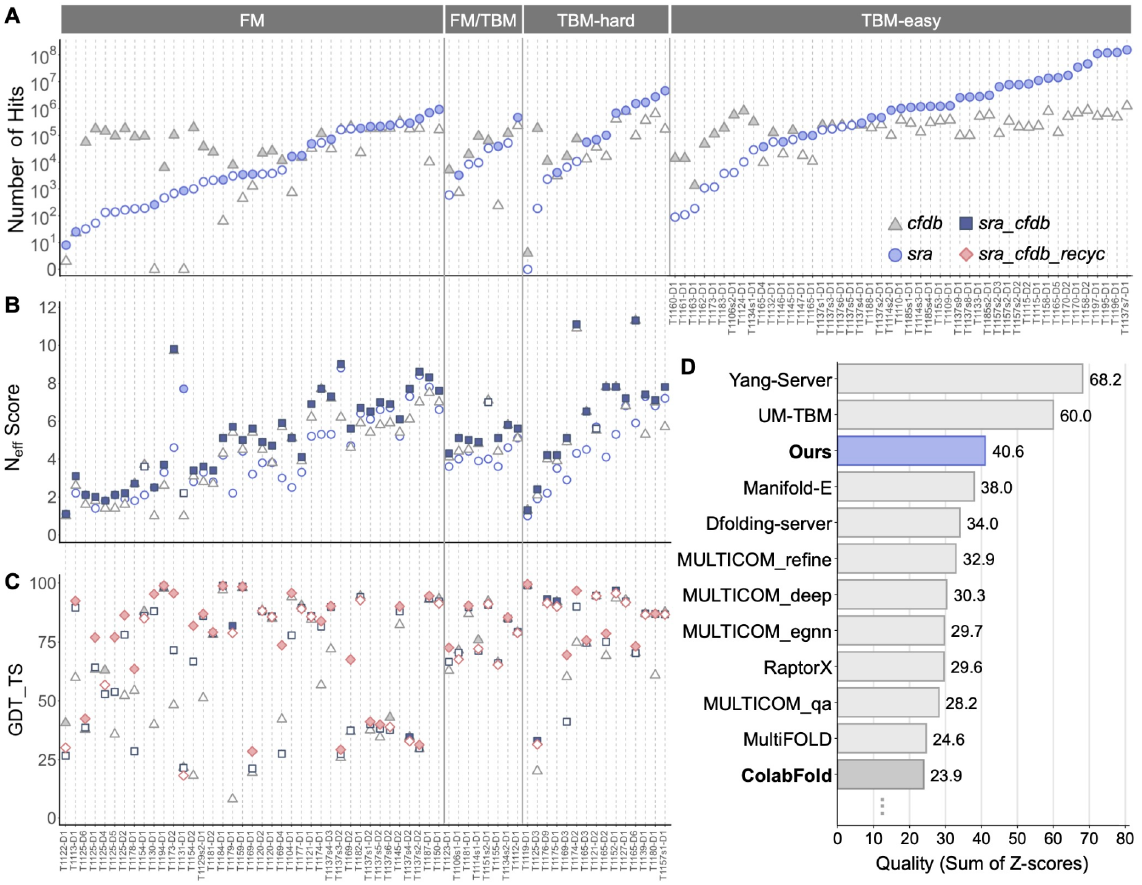
Effect of ColabFold parameters on structure prediction accuracy. **(A)** Comparison of homology search of 109 domains of 77 CASP15 targets. Each mark denotes the number of hits found for each target domain using either CF-search against CFDB (triangle) or MMseqs2 against SRA-mined and assembled proteins (circle) before the MSA filtering step. **(B)** N_eff_ scores of the different MSAs computed for each domain. The MSA with the most homologs and the highest N_eff_ is indicated with a filled mark in panels A and B, respectively. **(C)** Structure prediction of 62 target domains in the categories: FM, FM/TBM, and TBM-hard was evaluated based on GDT_TS scores of three prediction strategies: *cfdb* MSA, *sra_cfdb* MSA and *sra_cfdb_recyc*. The best-scoring strategy for each target domain is indicated with filled marks. **(D)** Prediction performance comparison between server groups in CASP15. The x-axis refers to the Sum Z (> 0.0) in Table 3. The score of this study is from the Model1 in Table 3. Here, ColabFold refers to the performance of the server group submitted in CASP15.

#### 1.2 Diving deeper

Serratus’ ability to scan the SRA in feasible time comes at the expense of its sensitivity to detect remote homologs. Specifically, it is limited in its ability to detect sequences with less than 50% identity to the query (Edgar et al. 2022). This prompted us to search deeply for remote homologs. To that end, we merged for each target its *cfdb* and *sra* MSAs (Fig. 1 ➂), resulting in an *sra_cfdb* MSA. We then ran HHblits with each *sra_cfdb* MSA as input against UniRef30 and BFD (Fig. 1 ➃), setting parameters to include all sequences in the output without any filtering and ensure maximum sensitivity (for a full parameter description, see Supp. information of Jumper et al. 2021). The resulting MSAs, denoted as *hh_sra_cfdb* MSAs, contained the input sequences as well as the homologs detected by HHblits.

To filter each input MSA based on the sequence identity (seqid) between its members and the query, we utilized a newly introduced filter module called filtera3m in MMseqs2, which implements ColabFold’s MSA filtering strategy. We removed unlikely homologs (seqid < 0.2), non-informative homologs (seqid > 0.95) and redundant sequences, keeping the most diverse and informative set of sequences in the MSA.

The filtered *cfdb* MSAs had on average 2,395 sequences, *sra_cfdb* MSAs -5,731 and *hh_sra_cfdb* MSAs - 8,133. We used HHmake (Steinegger et al. 2019a) to compute the number of effective sequences (N_eff_), where higher values indicate less similarity between the sequences and more diverse MSAs. Here, a more moderate increase was observed with the average N_eff_ score, rising from 4.87 for *cfdb* MSA to 5.43 for *sra_cfdb* MSA and to 7.26 for *hh_sra_cfdb* MSA (Fig. 2B).

### 2. The effect of homologs on structure prediction

In order to measure the effect of the various homolog collection strategies on structure prediction, we provided CF-predict with different input MSAs. These included the *sra_cfdb* and *hh_sra_cfdb* MSAs as well as their controls, which did not include Serratus-detected homologs from the SRA: *cfdb* and *hh_cfdb* MSAs. The *hh_cfdb* MSAs were produced in a similar manner to *hh_sra_cfdb* MSAs, using *cfdb* MSAs, rather than *sra_cfdb* MSAs, as the input to HHblits.

For each strategy, five models were produced and the best one was selected according to its computed predicted local distance difference test (pLDDT) score (Jumper et al. 2021). For the selected models, we measured the domain level accuracy, using the GDT_TS score (Zemla 2003). Compared to using *cfdb* MSAs, *sra_cfdb* MSAs significantly improved GDT_TS scores (Table 3), increasing scores for 38 out of 62 domains (Fig. 2C). On the other hand, using *hh_sra_cfdb* MSAs did not lead to a significant improvement over *sra_cfdb* MSAs (Table 3) and is therefore omitted from the figure.

**Table 3.** Effects of different strategies on protein structure prediction performance

| CFDB | SRA | HHblits | templates | multimer | 12 recycles | GDT_TS<sup>b</sup> | Sum Z (> 0.0)<sup>c</sup> |
| --- | --- | --- | --- | --- | --- | --- | --- |
| ✓ |  |  |  |  |  | 65.80 ± 25.58 | 17.09 |
| ✓ | ✓ |  |  |  |  | 70.97 ± 24.40<sup>d</sup> | 30.30 |
| ✓ |  | ✓ |  |  |  | 67.36 ± 25.15 | 15.82 |
| ✓ | ✓ | ✓ |  |  |  | 71.44 ± 23.30<sup>e</sup> | 26.32 |
| ✓ | ✓ |  | ✓ |  |  | 70.94 ± 23.99 | 31.60 |
| ✓ | ✓ |  |  | ✓ |  | 70.98 ± 24.12 | 30.97 |
| ✓ | ✓ |  |  |  | ✓ | 75.54 ± 22.17<sup>f</sup> | 40.59 |
| Model1<sup>a</sup> |  |  |  |  |  | 75.56 ± 22.12 | 40.56 |

<sup>a</sup> Model1 is the set of predicted-best strategies among all examined strategies, chosen based on pLDDT for each target domain.
<sup>b</sup> GDT\_TS scores are represented as mean ± standard deviation.
<sup>c</sup> Z-scores are calculated from each target domain's GDT\_TS, based on the server groups' mean and standard deviation. Sum Z (> 0.0) refers to the sum of positive Z-scores. All strategies were compared to their controls using the Wilcoxon signed-rank test. No significant improvement ( $p > 0.05$ ) was found except for:
<sup>d</sup> *sra\_cfdb* over *cfdb* MSAs ( $p = 0.0116$ )
<sup>e</sup> *hh\_sra\_cfdb* over *hh\_cfdb* MSAs ( $p = 0.0126$ )
<sup>f</sup> *sra\_cfdb\_recyc* over *sra\_cfdb* MSAs ( $p = 0.0003$ )

We further examined the relative performance of each strategy compared to other CASP15 servers using Z-scores, as follows. For each strategy, we deducted from its GDT_TS scores the mean servers’ GDT_TS score and divided it by the servers’ standard deviation. The sum of non-negative Z-scores and the average GDT_TS of all evaluated target domains were used as representative scores for each strategy (Table 3). As expected, the addition of *sra* to *cfdb* MSAs substantially improved the performance, increasing the sum of Z-scores from 17.09 to 30.30 (Table 3). On average, no significant improvement was observed when adding HHblits-detected homologs (Table 3). However, specific domains gained a substantial improvement by including these homologs, indicating a variable effect for each target. A notable example with the highest improvement is target T1178-D1, where running HHblits increased GDT_TS from 28.61 for *sra_cfdb* to 84.17 for *hh_sra_cfdb* MSAs.

### 3. Tuning parameters

In addition to homolog collection strategies, we examined the impact of three advanced features of CF-predict: using templates, using multimer models, and increasing the number of recycles. The strategies corresponding to these features were built on the MSAs constructed in previous steps and are denoted: *sra_cfdb_temp, sra_cfdb_recyc* and *sra_cfdb_mult* and their control: *sra_cfdb*. Other strategies, taken by the leading CASP15 servers are detailed in Table 4.

**Table 4.**
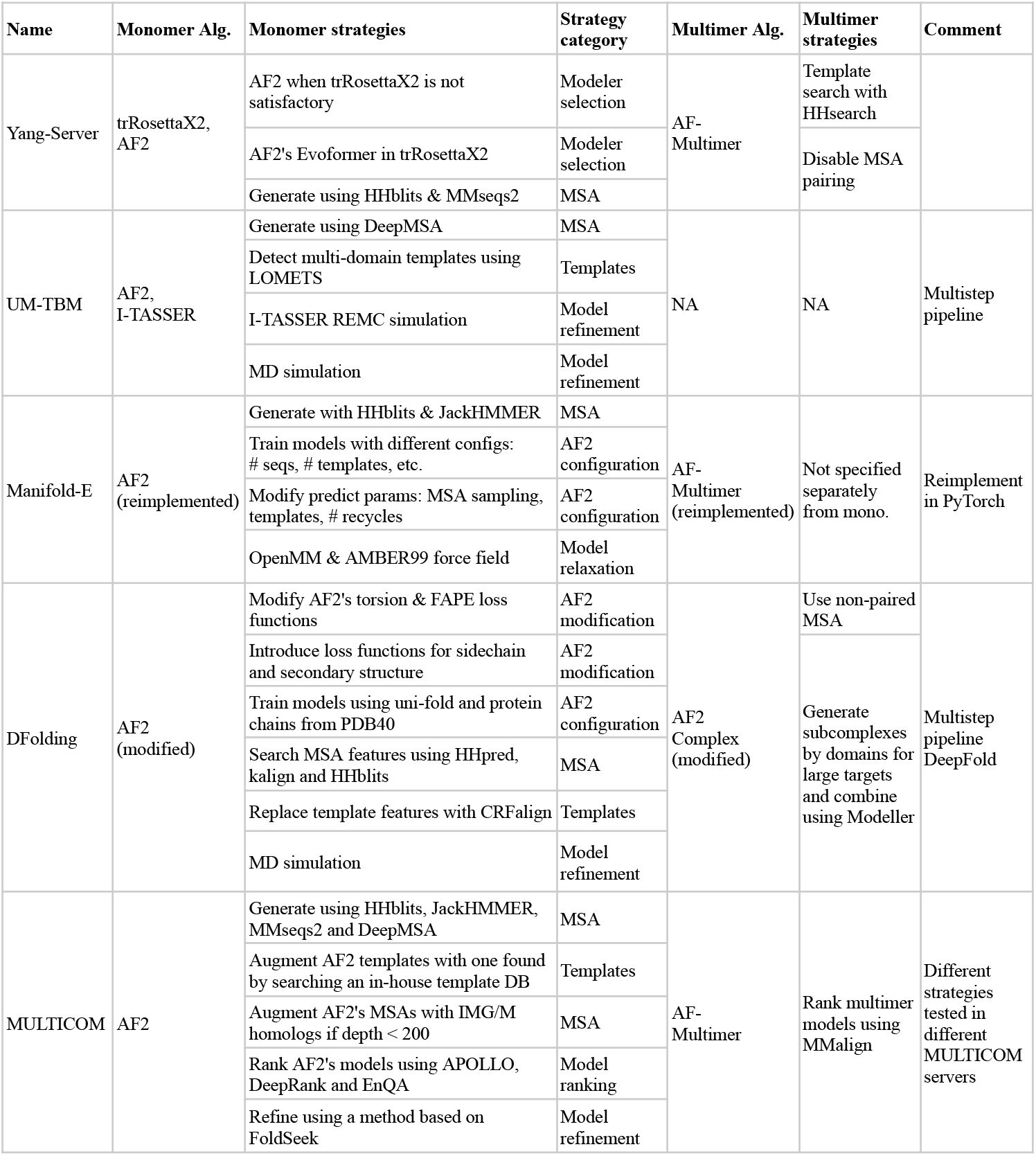
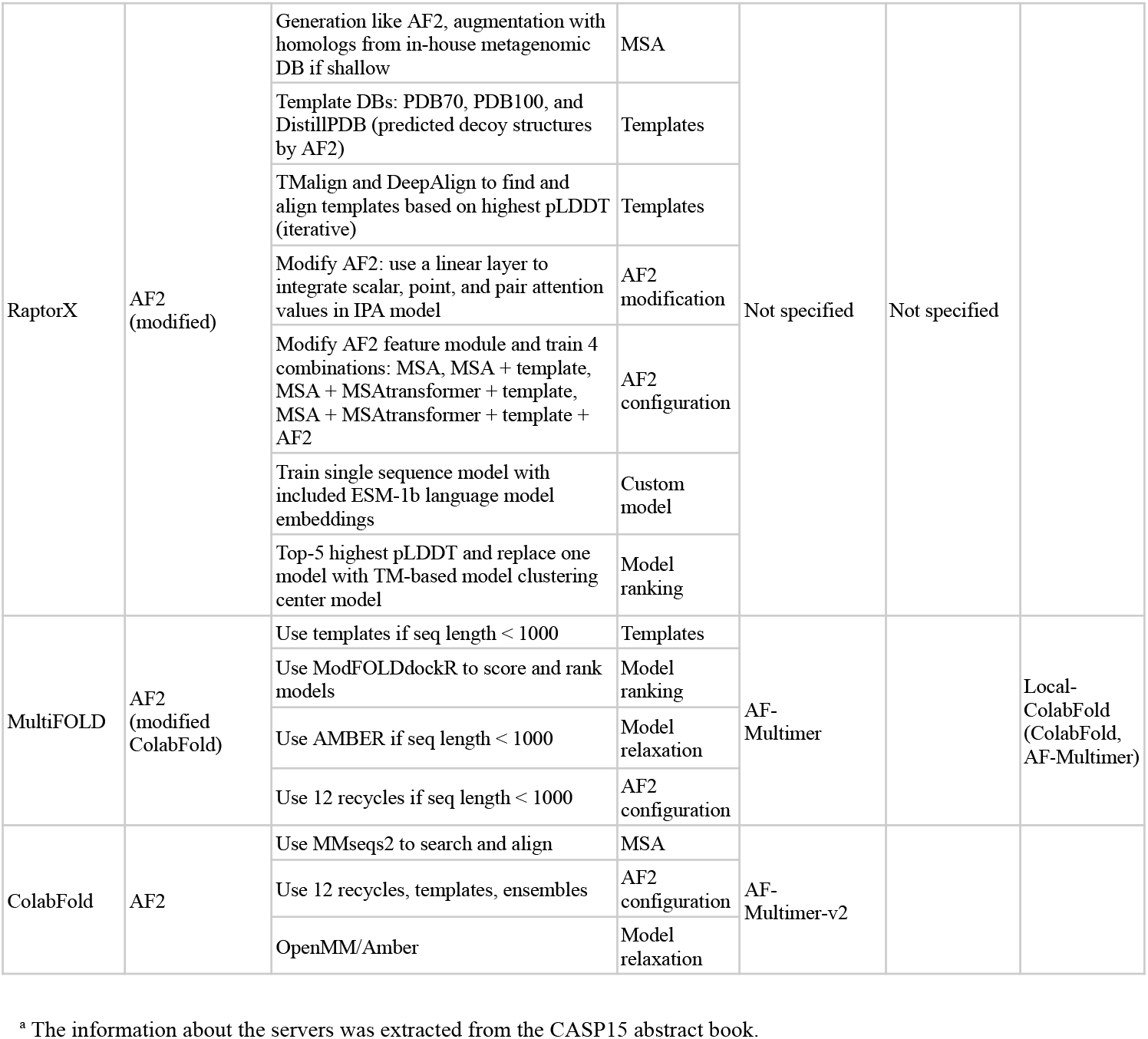
Use of prediction algorithms and strategies among leading CASP15 servers^a^.

#### 3.1 Leveraging templates

Two out of the five AlphaFold2 models require structural features as input. Setting the “templates” parameter (Fig. 1 ➄) changes the default behavior of CF-predict from using mock templates to querying the PDB70 (Steinegger et al. 2019a) using the UniRef30-based constructed profile as input and returning hits, which are later aligned by HHsearch.

Adding templates did not improve the accuracy compared to using default parameters (Table 3). However, it should be noted that our examined targets are FM, FM/TBM and TBM-hard, which are classified as difficult targets to find templates for. Applying templates to TBM-easy targets might have different effects on prediction performance.

#### 3.2. Multimer modeling

We also tested the impact of setting on the “multimer” option (Fig. 1 ➅). This changes the default behavior of CF-predict from treating the input query sequence as a monomer to considering it to be a part of a complex. This has the potential of stabilizing the structure, thereby improving prediction accuracy. We applied multimer modeling only for the homo-oligomer targets, based on the stoichiometry provided by CASP and used the multimer model weights from AlphaFold-multimer version 2 (Evans et al. 2022).

Using multimer modeling had a diverse range of effects. Overall, it did not make a significant improvement over *sra_cfdb* MSAs (Table 3). However, it drastically increased or decreased prediction performance for certain targets. Two notable examples stand out: T1178-D1 achieved the highest improvement, with its GDT_TS score soaring from 28.61 to 61.02 after incorporating multimer modeling. Conversely, the GDT_TS score of target T1174-D1 experienced the largest decrease, dropping from 81.48 to 59.84.

#### 3.3 Adding more recycles

Through the “recycle” parameter (Fig. 1 ➆), CF-predict allows setting the number of iterations in which a prediction will be re-fed to the AlphaFold2 models. By default, this value is set to 3, but additional recycles have the potential to improve prediction accuracy (Mirdita et al. 2022). Thus, when exploring this option, we set it to 12. The recently released version 3 of AlphaFold-multimer (Evans et al. 2022) uses up to 20 recycle iterations, with early stopping if a model has already converged.

Increasing recycles significantly improved the prediction accuracy compared to the control *sra_cfdb* with default parameters (Table 3). As depicted in Figure 2C, *sra_cfdb_recyc* MSA scored higher than *cfdb* MSAs and *sra_cfdb* MSAs in 34 domains and achieved high-accuracy (GDT_TS > 70) in 72% of the 62 domains.

### 4. Strategy selection and comparison with CASP15 servers

Finally, among the seven examined strategies, we selected the one with the highest pLDDT for each target domain, denoted here as Model1. We then compared the performance of Model1 with the leading server groups in CASP15, including the original ColabFold server, which is similar to this study’s *cfdb* MSAs (Fig. 2D). Model1 resulted in an average GDT_TS of 75.56 and sum of Z-scores of 40.56, increasing 9.76 and 23.47 units from the baseline *cfdb* MSAs, respectively. Notably, 46 out of 62 domains (74%) of Model1 achieved high-accuracy scores (GDT_TS > 70), compared to 52% of the *cfdb* MSAs. When comparing with other server groups based on the sum of Z-scores, Model1 would have ranked 3rd, outperforming the ColabFold original server, which ranked 11th in CASP15 among server-only groups on non-easy targets.

In order to examine the validity of using pLDDT as selection criterion, we compared for each domain the GDT_TS score of Model1 and the highest GDT_TS score (Model_best), across all strategies. In case of perfect agreement between pLLDT and GDT_TS, these values should be equal. However, we observed a disagreement for 37 out of 62 target domains, resulting in a notable increase in Model_best with the average GDT_TS reaching 78.13 and the sum of Z-scores reaching 52.31. This disparity between pLDDT and GDT_TS highlights the challenge in selecting the best model. For instance, choosing the strategy with the highest pLDDT for target T1104-D1 yielded a GDT_TS score of 77.56 (*hh_sra_cfdb* MSA), while the actual best GDT_TS score was 95.73 (*sra_cfdb_recyc* MSA).

To address this discrepancy, there were attempts to use alternative model selection (ranking) methods in CASP15, instead of relying solely on pLDDT (Table 4). For instance, the MULTICOM servers (Liu et al. 2023) utilized APOLLO (Wang et al. 2011), DeepRank (Renaud et al. 2021), and EnQA (Chen et al. 2023a) for ranking models, and MultiFOLD (McGuffin et al. 2023) employed ModFOLDdockR (Edmunds et al. 2023) for both scoring and ranking purposes. Further developments of ranking methods are needed to improve the accuracy and reliability of model selection for structure prediction.

## Concluding remarks

In this study, we have shown the importance of a comprehensive inclusion of metagenomic sequences from the SRA for improving protein structure prediction.

Our results highlight the large variation in the number of homologs found for different targets. For example, over 100 million environmental homologs were found for T1137s7, T1195, T1196, and T1197 -the same order of magnitude as the entire UniProt database. However, there are some targets for which few matches were detected, possibly due to the limited sensitivity of the mining procedure.

Serratus, the tool used for mining the SRA has impressive capabilities, but also significant constraints. It is limited to detecting homologs, which have about 50% sequence identity to the query, missing the full potential of homologs from the twilight zone (Rost 1999). Fast and more sensitive search methods are thus required to further improve our ability to exploit the SRA. Additionally, using Serratus in a similar manner to this study is likely to cost thousands of dollars (Edgar et al. 2022) and this cost could become limiting with the expected continued exponential growth of the SRA.

We further investigated the impact of advanced CF-predict features on structure prediction performance. While adding more recycles led to improvement, using multimer models and templates did not contribute significantly. Nonetheless, each feature may have varying effects on different targets, as demonstrated by some notable examples. Thus, it is highly recommendable to experiment with different combinations of these features to optimize performance.

In conclusion, while limitations persist, advancements in metagenomic data mining tools, coupled with a blend of automated and human-guided predictions, promise exciting prospects for the future. Further, the results of this research underline the necessity for diversification in methodologies used in protein structure prediction. Finally, the disparity in MSA coverage among different targets stresses the importance of individual target evaluation and tailored approaches. As the field continues to evolve, we anticipate these findings to contribute to the ongoing quest for accurate protein structure prediction.

## Data availability

The data that support the findings of this study are available at https://doi.org/10.5281/zenodo.8126538.

## Acknowledgments

MS acknowledges the support by the National Research Foundation of Korea, grants [2020M3-A9G7-103933, 2021-R1C1-C102065, 2021-M3A9-I4021220], Samsung DS research fund and the Creative-Pioneering Researchers Program through Seoul National University. MM acknowledges support by the National Research Foundation of Korea (grant RS-2023-00250470). Computing resources were provided by the University of British Columbia Community Health and Wellbeing Cloud Innovation Centre, powered by AWS. RC was supported by ANR grants [ANR-19-CE45-0008, ANR-22-CE45-0007, PIA/ANR16-CONV-0005, ANR-19-P3IA-0001], and H2020 Marie Skłodowska-Curie grants [No 956229 and 872539]. AK acknowledges the support of the US National Institute of General Medical Sciences (NIGMS/NIH), grant R01GM100482.

## Notes

https://doi.org/10.5281/zenodo.8126538

### Competing Interest Statement

The authors have declared no competing interest.

